## Supplementary figures and images for "Mixed culture metagenomics of the microbes making sour beer"

### Supplementary Information File S2

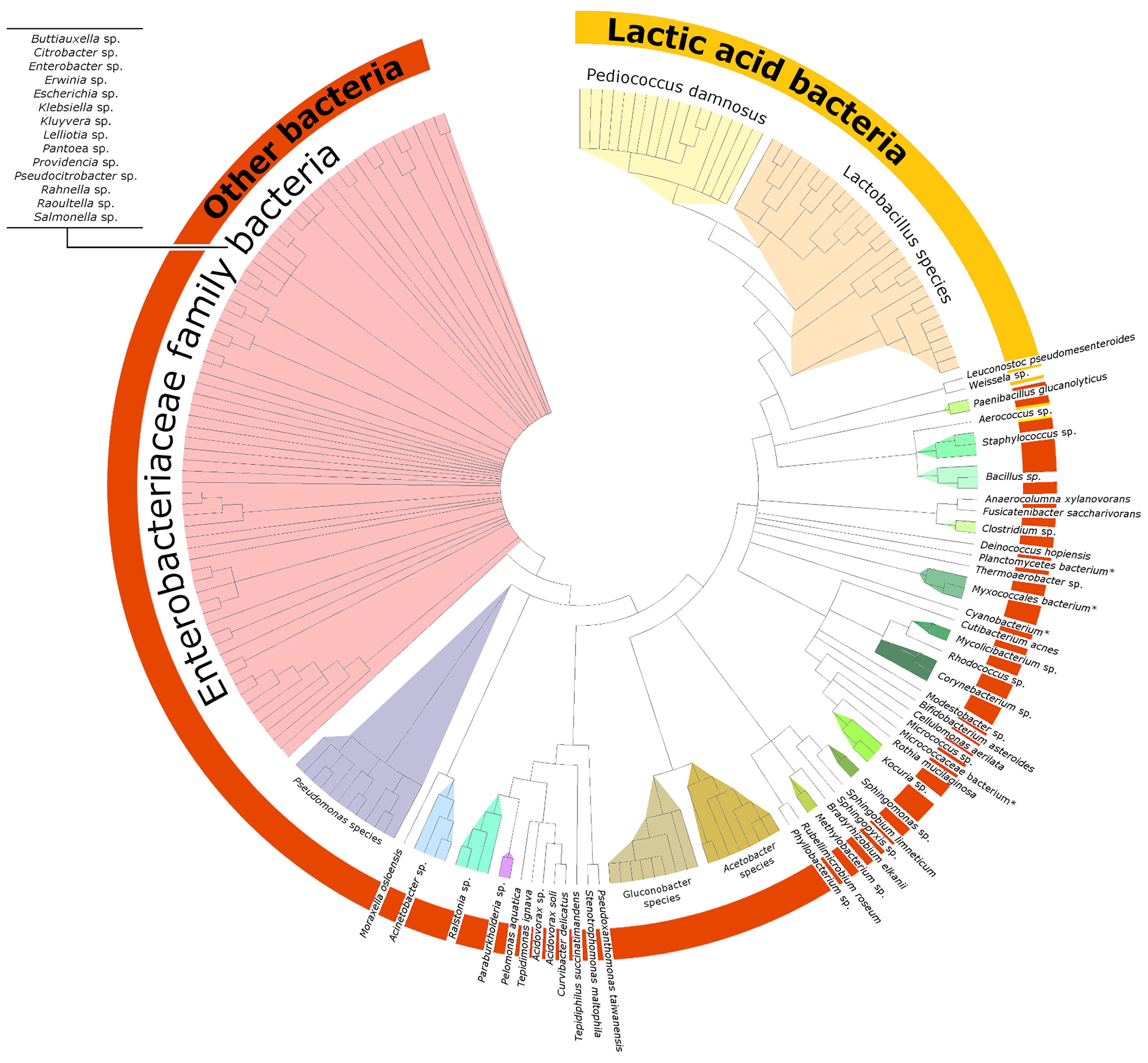

### Supplementary Information File S3

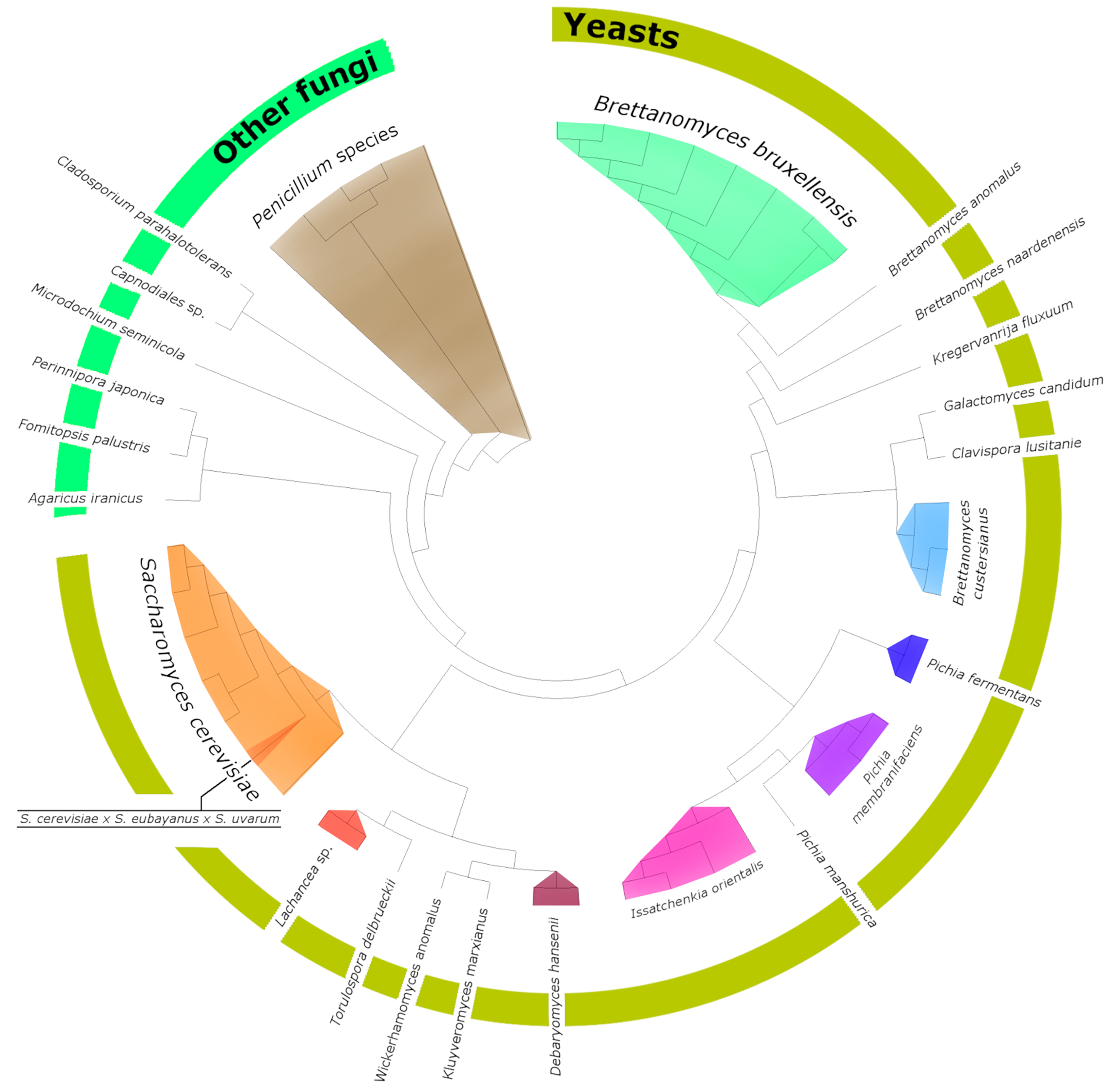
