## Supplementary Table S1 for "Mixed culture metagenomics of the microbes making sour beer"

| **Sample** | **Commercial culture inoculated*** | **Identification match** |
| --- | --- | --- |
| #2 | New World Saison Blend Escarpment Labs (*Saccharomyces* spp. + *Brettanomyces* spp.)  Brett M Escarpment Labs (*Brettanomyces bruxellensis*) | *Saccharomyces cerevisiae* was identified in 61.4% of ASVs from fungi  *Brettanomyces bruxellensis* was identified in 30.3% of ASVs from fungi |
| #11 | Sour Solera Bootleg (*Saccharomyces* spp. + *Brettanomyces* spp. + *Lactobacillus* spp. + *Pediococcus* spp. + other wild microorganisms  Mélange Yeast Bay (Saccharomyces cererevisiae var. diastaticus + *Saccharomyces fermentati* + *Brettanomyces* spp. + *Lactobacillus brevis* + *Lactobacillus delbrueckii* + *Pediococcus damnosu*s)  Bugcounty East Coast Yeast (*Saccharomyces* spp. + *Brettanomyces lambicus* + *Brettanomyces anomalus* + *Brettanomyces clausseni* + *Brettanomyces custersianus* + *Brettanomyces naardenensis* + *Lactobacillus* spp. + *Pediococcus* spp.) | *Saccharomyces cerevisiae* was identified in 77.9% of ASVs from fungi  *Brettanomyces bruxellensis* in 13.6% of ASVs from fungi  *Brettanomyces* *anomalus* in 6.7% of ASVs from fungi  *Brettanomyces* *naardenensis* in 0.8% of ASVs from fungi  *Brettanomyces* *custersianus* in 0.3% of ASVs from fungi  *Lactobacillus* *casei-paracasei* in 74.0% of ASVs from bacteria  *Pediococcus* *damnosus* in 13.8% of ASVs from bacteria  *Lactobacillus* *brevis* in 9.9% of ASVs from bacteria  *Lactobacillus* *acetotolerans* in 0.5% of ASVs from bacteria  *Lactobacillus* *plantarum* in 0.2% of ASVs from bacteria  *Lactobacillus* *delbrueckii* in 0.06% of ASVs from bacteria |
| #18 | Roeselare blend Wyeast (*Brettanomyces* spp. + *Saccharomyces cerevisiae* + *Lactobacillus* spp. + *Pediococcu*s spp.)  French Saison Wyeast (*Saccharomyces cerevisiae* var. *diastaticus*)  WLP650 White Labs (*Brettanomyces bruxellensis*) | *Brettanomyces* *bruxellensis* was identified in 99.5% of ASVs from fungi  *Saccharomyces* *cerevisiae* was identified in 0.09% of ASVs from fungi  *Lactobacillus* *acetotolerans* was identified in 100% of ASVs from bacteria |
| #19 | Belgian Saison I White Labs (*Saccharomyces cerevisiae*) | *Saccharomyces* *cerevisiae* was identified in 7.3% of ASVs from fungi |

**Supplementary Table S1:** Commercial cultures used to inoculate non-spontaneous mixed-fermentation beers and matches found in the next-generation sequencing data.

*The composition of the commercial culture was obtained from company/laboratory website.
